# Spatial Heterogeneity and Subtypes of Functional Connectivity Development in Youth

**DOI:** 10.1101/2025.01.24.634828

**Authors:** Hongming Li, Zaixu Cui, Matthew Cieslak, Taylor Salo, Tyler M. Moore, Raquel E. Gur, Ruben C. Gur, Russell T. Shinohara, Desmond J. Oathes, Christos Davatzikos, Theodore D. Satterthwaite, Yong Fan

## Abstract

The brain functional connectome development is fundamental to neurocognitive growth in youth. While brain age prediction has been widely used to assess connectome development at the individual level, traditional approaches providing a global index overlook the spatial variability and inter-individual heterogeneity of functional connectivity (FC) development across the cortex. In this study, we introduced a regional brain development index to assess spatially fine-grained FC development. We examined the spatial variability of FC development and stratified individuals into subtypes with distinct patterns of spatial heterogeneity in region-wise FC development across the cortex through clustering. An evaluation conducted on a sample of youths aged 8-23 years using fMRI data from the Philadelphia Neurodevelopmental Cohort (PNC) revealed three distinct FC development subtypes. Individuals with advanced FC development aligned with the hierarchical brain organization along the sensorimotor-association (S-A) axis demonstrated superior cognitive performance compared to those with other patterns. These patterns were replicated in the Human Connectome Project Development (HCP-D) cohort, confirming their robustness. Further analysis revealed associations between FC development and gene expression, with enriched genes linked to neural differentiation, synaptogenesis, and myelination. These findings suggest that spatial heterogeneity in FC development reflects underlying cortical microstructure and hierarchical cortical organization, underscoring its critical role in understanding neurocognitive maturation and individual variability during youth.

## Introduction

The development of functional networks is a dynamic and complex process closely linked to ongoing maturation of the brain’s functional organization, which underpins the progressive enhancement of cognitive, emotional, and social capacities throughout childhood and adolescence [1, 2]. Recent studies have shown that the spatial-temporal patterns of functional network development in youth systematically align with cortical hierarchy organization, particularly along the sensorimotor-association (S-A) axis, with developmental increases in functional connectivity (FC) in sensorimotor cortices and decreases in association cortices [3]. Neurocognition undergoes a protracted development trajectory parallel to the maturation of the brain’s functional organization, with individual differences in youth linked to important outcomes such as academic performance and quality of life [2], while deficits in neurocognition are associated with vulnerability to mental illness. Therefore, advancing our understanding of FC development at the individual level may provide valuable insights into typical development processes and deviations from these typical trajectories in the context of mental illness.

Brain age has increasingly been utilized to evaluate brain status and track brain changes at the individual level, leveraging neuroimaging data across various modalities [4-19]. The difference between brain age and chronological age, known as the brain age gap (BAG), provides a quantitative and interpretable metric for characterizing brain changes associated with development, aging, and brain disorders. BAG measures derived from functional connectome have revealed that individuals with accelerated brain development exhibit enhanced FC in specific functional brain networks, including the default mode network (DMN), during childhood and adolescence [9]. Existing studies have established a linkage between brain age derived from structural imaging data and cognition, demonstrating that individuals with advanced brain development exhibit superior cognitive processing speed compared to those with delayed development [5]. While studies have identified associations between FC and neurocognition in youth using predictive modeling [20-23], it remains unclear whether and how brain age estimates derived from the functional connectome are associated with neurocognition in youth.

Most existing studies estimate brain age from a global perspective by mapping whole-brain, high-dimensional FC profiles to a single unitary index, which inherently overlooks the spatial variability of FC changes across brain regions. To address this limitation, several studies have proposed fine-grained modeling at the regional level [7, 10, 12, 14, 15, 17, 24]. Distinct and heritable patterns of brain aging, derived from structural imaging features, have been linked to brain disorders, such as schizophrenia, multiple sclerosis, and dementia [7]. Additionally, heterogenous aging patterns across brain regions in schizophrenia have been revealed using multimodal neuroimaging data [17]. Advanced brain aging in specific brain regions has also been shown to mediate cognitive outcomes in cerebral small vessel disease, highlighting the potential to capture heterogeneous brain changes beyond a unitary measure of global brain age [12]. Given that variations in FC profiles, functional topography, and structural-functional coupling are associated with brain development and neurocognition [2, 3, 6, 8, 9, 20, 21, 23-26], exploring regional patterns of brain development measures derived from FC and their associations with neurocognition could provide valuable insights into the development of brain functional organization and neurocognitive processes.

In this study, we utilized a large sample (total *n*=1149) of youths aged 5-23 years from the Philadelphia Neurodevelopmental Cohort (PNC) [27] and the Human Connectome Project Development cohort (HCP-D) [28] to investigate region-wise FC development using functional MR imaging (fMRI) data. For each cortical region, we developed a region-specific age prediction model based on its regional FC profile. Each model provides an individualized brain age measure, and the difference between this measure and chronological age, referred to as regional brain development (RBD) index, reflects region-wise FC development. We hypothesized that the RBD index would exhibit spatial variability and inter-subject heterogeneity, linked to developmental differences in neurocognition. To explore the spatial variability of regional FC development and its associations with neurocognition, we stratified individuals into subtypes characterized by distinct FC development patterns using a cluster analysis based on individual subjects’ whole-brain RBD indices. Subsequently, we investigated differences in cognitive measures across these subtypes. Our analysis revealed that individuals with advanced FC development parallel to the hierarchical cortical organization defined by the sensorimotor-association (S-A) axis exhibited superior cognitive profiles. To investigate the biological mechanism underlying the spatial variability of FC development, we examined the relationship between regional FC development patterns and gene expression. We predicted that regional FC development patterns aligning with the cortical hierarchy organization would show enrichment in the expression of genes associated with neuron development.

## Results

We used fMRI data from the PNC (*n*=693; ages 8-23) to examine the spatial variability and inter-individual heterogeneity in regional FC development. The RBD index, defined as brain age minus chronological age and corrected for bias and covariates, was used as a surrogate measure to assess FC development across cortical regions. Specifically, for each cortical region, a dedicated brain age prediction model was built on its FC profile, which was defined as its FC measures to all the other cortical regions, with Pearson’s correlation coefficient serving as the FC measure. Five-fold cross-validation was implemented to predict brain age across all individuals and cortical regions. An overview of the analysis process is illustrated in Fig. 1. Using the RBD index, we first investigated spatial variability in FC development and its inter-individual heterogeneity. Next, we identified subgroups of individuals with distinct patterns of regional FC development and explored their associations with neurocognitive development. To deepen our understanding, we examined how distinct patterns of FC development align with fundamental properties of functional and structural brain organization. Finally, we assessed the relationship between these FC development patterns and gene expression to explore the underlying biological mechanism.

**Fig 1.**
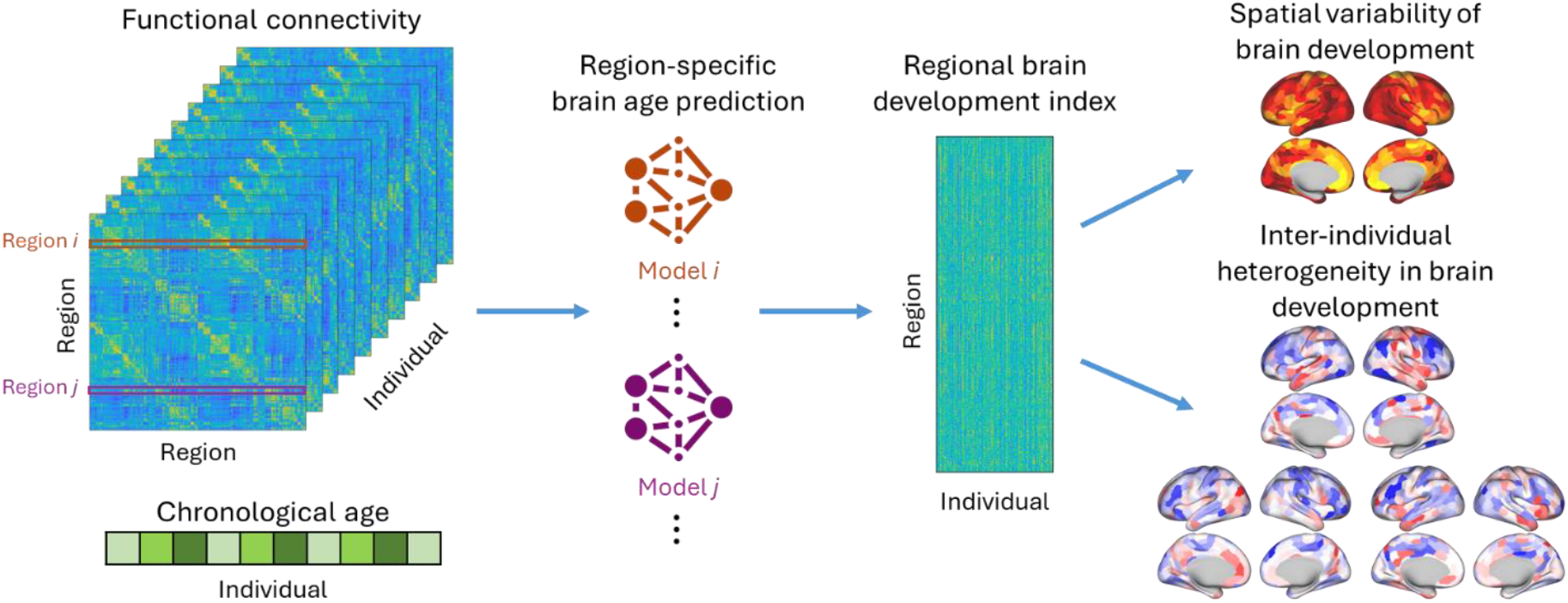
The regional brain development (RBD) index derived from functional connectivity (FC). Region-specific brain age prediction models were built upon regional FC profiles comprising FC measures of each region with all other cortical regions, to assess region-wise FC development. These prediction models were trained using a 5-fold cross-validation approach to predict brain age for each individual and cortical region. Subsequently, the spatial variability in prediction performance and inter-individual heterogeneity in the RBD index were analyzed to provide deeper insights into region-wise FC development.

### Spatial variation and inter-individual heterogeneity in functional connectivity development

To characterize the spatial variation of FC development across cortical regions, we analyzed differences in their capacity for age prediction. Two metrics, accuracy (*r*) and mean absolute error (MAE), were used to quantify each region’s predictive capacity, with accuracy (*r*) computed as the Pearson’s correlation coefficient between the predicted age (prior to bias correction) and chronological age. Cortical regions exhibited varying capacity for age prediction (Fig. 2A), with certain functional networks showing higher predictive capacity, as indicated by higher *r* and lower MAE values. These networks included the somatomotor network, ventral attention network (VAN), and frontoparietal network (Fig. 2B). However, heterogeneity was evident within individual functional networks: for example, the medial prefrontal aspect of the DMN displayed higher predictive capacity compared with lateral elements of the DMN.

**Fig 2.**
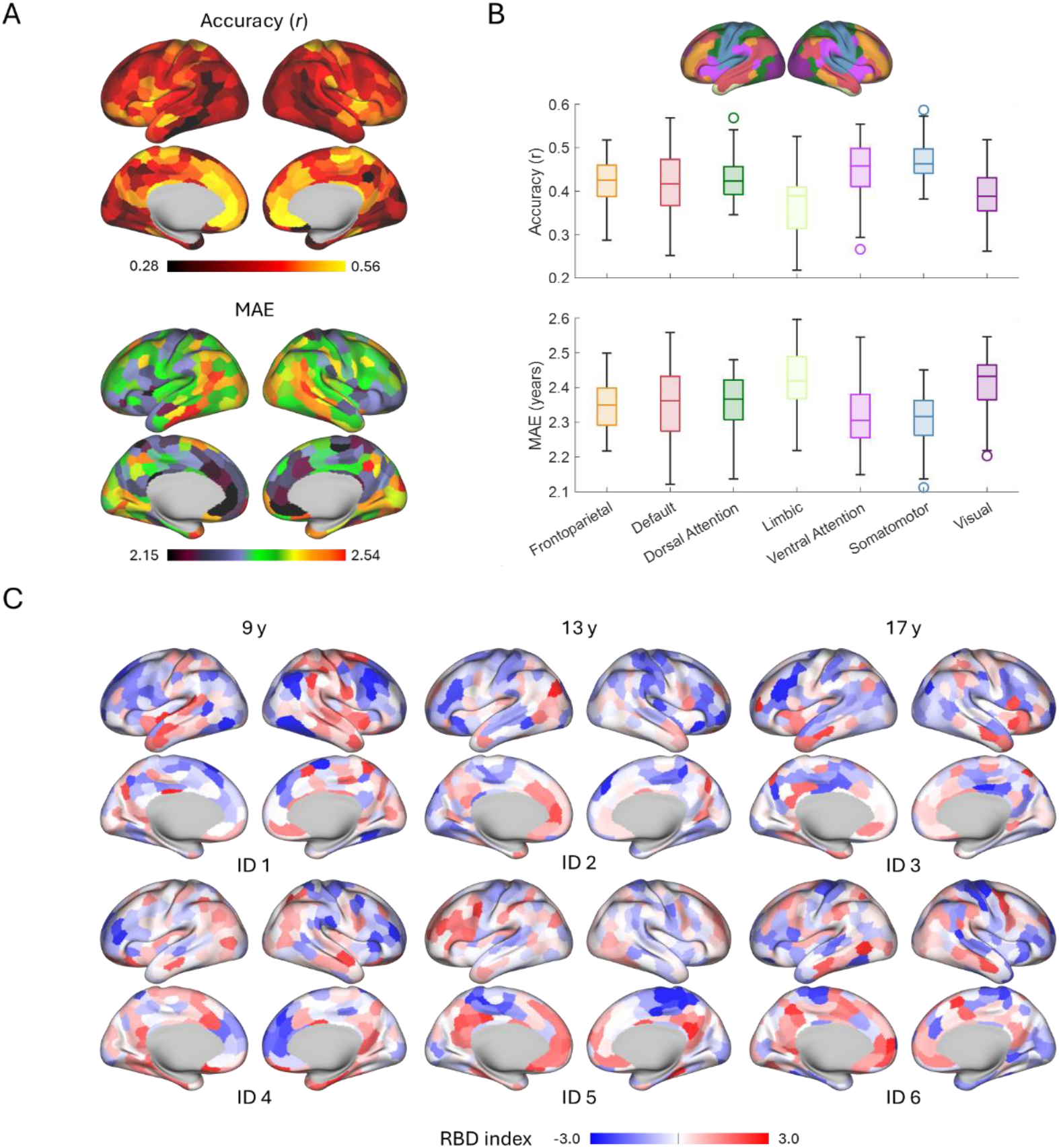
Spatial variations and inter-individual differences in functional connectivity development. (A) Regional capacity for brain age prediction, quantified by accuracy (*r*) and mean absolute error (MAE), reflects spatially varying FC development. (B) Distribution of region-wise age prediction capacity across brain functional networks based on the Yeo 7-network atlas [29]. (C) Maps of RBD indices are shown for six randomly selected individuals (IDs 1-6) at three developmental time points: ages 9, 13, and 17 (two individuals per time point), illustrating brain regions with advanced (positive, red) or delayed (negative, blue) FC development.

Building on the observed variation in FC development across cortical regions, we next investigated inter-individual differences in the pace of regional FC development. To assess this, we quantified the brain development for each cortical region and individual using the RBD index, defined as brain age minus chronological age and corrected for bias and covariates [30], with positive/negative values indicating advanced/delayed development. Individuals of the same chronological age (9, 13, or 17 years) showed variability in region-wise brain development (Fig. 2C), highlighting inter-individual variability in the pace of regional FC development. Additionally, distinct spatial covariation patterns in regional brain development were evident at the individual level. For instance, the 13-year-old individual (Fig. 2C, bottom-middle) exhibited consistently advanced development in fronto-parietal and DMN regions alongside delayed development in sensorimotor regions, whereas the 9-year-old individual (Fig.2C, top left) showed a largely opposite pattern.

### Subtypes of functional connectivity development and their links to neurocognition

To investigate the observed inter-individual heterogeneity and potential spatial covariation in regional FC development, we identified covarying patterns of FC development and stratified individuals into subgroups based on their whole-brain RBD index maps. Using the K-means clustering algorithm, individuals were grouped into three subtypes, with each subtype characterized by a distinct FC development pattern (P1, P2, and P3) represented by the centroids of their RBD index maps (Fig. 3A). Specifically, P1 showed a global trend of delayed FC development across cortical regions; P2 exhibited advanced FC development localized to the association cortex, prominently within DMN regions; P3 displayed widespread advanced FC development, particularly pronounced in sensorimotor regions.

**Fig 3.**
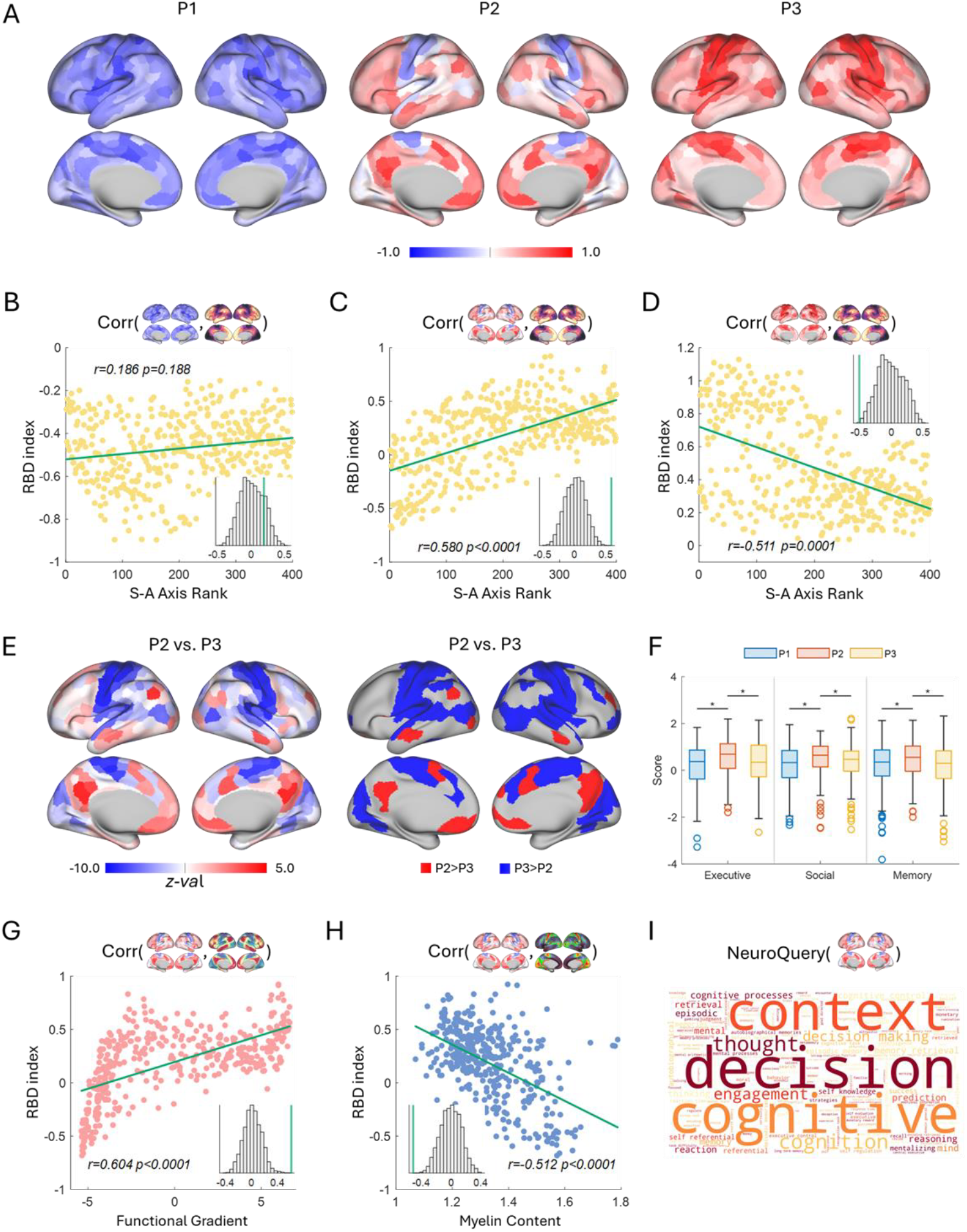
Subtypes of functional connectivity development identified based on RBD index maps. (A) Centroid RBD index maps of the three subgroups identified in the PNC cohort. (B-D) Alignment of the development patterns with the sensorimotor-association (S-A) axis rank across cortical regions, with each dot representing one cortical region. (E) Region-wise comparison of RBD indices between P2 and P3, with statistical significance determined using the Wilcoxon rank sum test (*p*<0.05, FDR-corrected). (F) Statistical comparisons of cognition scores across three subgroups, highlighting differences in cognitive performance. (G) Significant correlation between the development pattern of P2 and the principal functional gradient (*p*<0.0001, Spin-test). (H) Significant negative correlation between the development pattern of P2 and the cortical myelin content (*p*<0.0001, Spin-test). (I) Meta-analytic functional decoding result of the development pattern of P2 based on the NeuroQuery, where larger font size indicates stronger correlation with specific functional term.

To further characterize the spatial variation of these distinctive development patterns, we examined their alignment with the S-A axis, which represents a unified cortical organization across diverse neurobiological properties and has been demonstrated to be significantly associated with FC development [3]. While the development pattern of P1 did not significantly align with the S-A axis (Fig. 3B, Spin-test, *r*=0.186, *p*=0.188), those of P2 and P3 showed significant correlations, with P2 showing a strong positive correlation (Fig. 3C, Spin-test, *r*=0.580, *p*<0.0001) and P3 showing a strong negative correlation (Fig. 3D, Spin-test, *r*=-0.511, *p*=0.0001). Notably, the cortical regions with predominantly advanced development in the P2 were located at the high-ranking association end of the S-A axis, while those in P3 were located at the low-ranking sensorimotor end. Given the overall advanced FC development of P2 and P3, we then examined their differences in FC development across regions. Specifically, P2 demonstrated significantly more advanced FC development in DMN regions compared with P3, whereas P3 demonstrated significantly greater advanced development in sensorimotor regions relative to P2 (Fig. 3E).

We next analyzed the cognitive performance of individuals across subtypes in three cognitive domains: executive function, social cognition, and memory, to investigate their behavioral implications. Individuals in P2 demonstrated significantly higher cognitive performance than those in P1 and P3 (Wilcoxon rank sum test, *p*<0.05, Fig. 3F). Interestingly, no significant differences were observed between P1 and P3, despite globally advanced FC development of individuals in P3 (Fig. S1B). There were no significant differences across subgroups regarding chronological age and sex. These findings highlight the critical role of spatial heterogeneity in FC development during youth, particularly within regions that support neurocognitive functions.

To better understand the relationship between distinctive FC development patterns and properties of the brain connectome, we also analyzed their alignment with the principal functional gradient and cortical myelin content. The P2 showed a strong positive correlation with the principal functional gradient (Fig. 3G, Spin-test, *r*=0.604, *p*<0.0001), consistent with prior findings that hierarchical functional organization supports neurocognition development. Moreover, the P2 exhibited a significantly negative correlation with cortical myelin content (Fig. 3H, Spin-test, *r*=-0.512, *p*<0.0001), suggesting a relationship with the ongoing process of myelination during brain maturation. On the other hand, the P3 exhibited a reversed alignment pattern, showing a negative correlation with the principal functional gradient and a positive correlation with the cortical myelin content (Fig. S2). These findings underscore the critical interplay between functional organization and structural maturation of brain connectome in shaping neurocognitive outcomes.

In addition, we utilized a meta-analytic functional decoding technique [31] based on the NeuroQuery database [32] to interpret the functional significance of the identified FC development patterns. The decoding results also revealed significant associations between the P2 and neurocognition related terms such as cognitive, decision, and context (Fig. 3I).

### Associations between regional FC development and gene expression

To investigate the biological mechanism underlying the regional FC development, we analyzed the relationship between the FC development patterns and gene expression data from the Allen Human Brain Atlas (AHBA) [33]. We first examined the alignment of FC development patterns with the first principal component (PC1) of gene expression [34]. The P2 exhibited significant negative correlation (Fig. 4A, Spin-test, *r*=-0.440, *p=*0.0017), while no significant correlations were observed for P1 and P3 (Fig. S3). We then used spatial correlation analysis to examine the association between the P2 pattern and the expression of 4,112 genes in the cortex. This analysis identified 735 positively correlated genes and 783 negatively correlated genes (*q*<0.05, FDR corrected, Spin-test). Among them, the neuronal differentiation 6 (NEUROD6) gene was the most positively correlated, while the Mitochondrial Ribosomal Protein L48 (MRPL48) gene was the most negatively correlated (Fig. 4B). Using Specific Expression Analysis (SEA) [35], we found that the set of positively correlated genes was enriched for neurodevelopmental genes with cortical expression spanning stages from early childhood to young adulthood (Fig. 4C), suggesting that these genes may play a role in supporting the regional FC development in youth.

**Fig 4.**
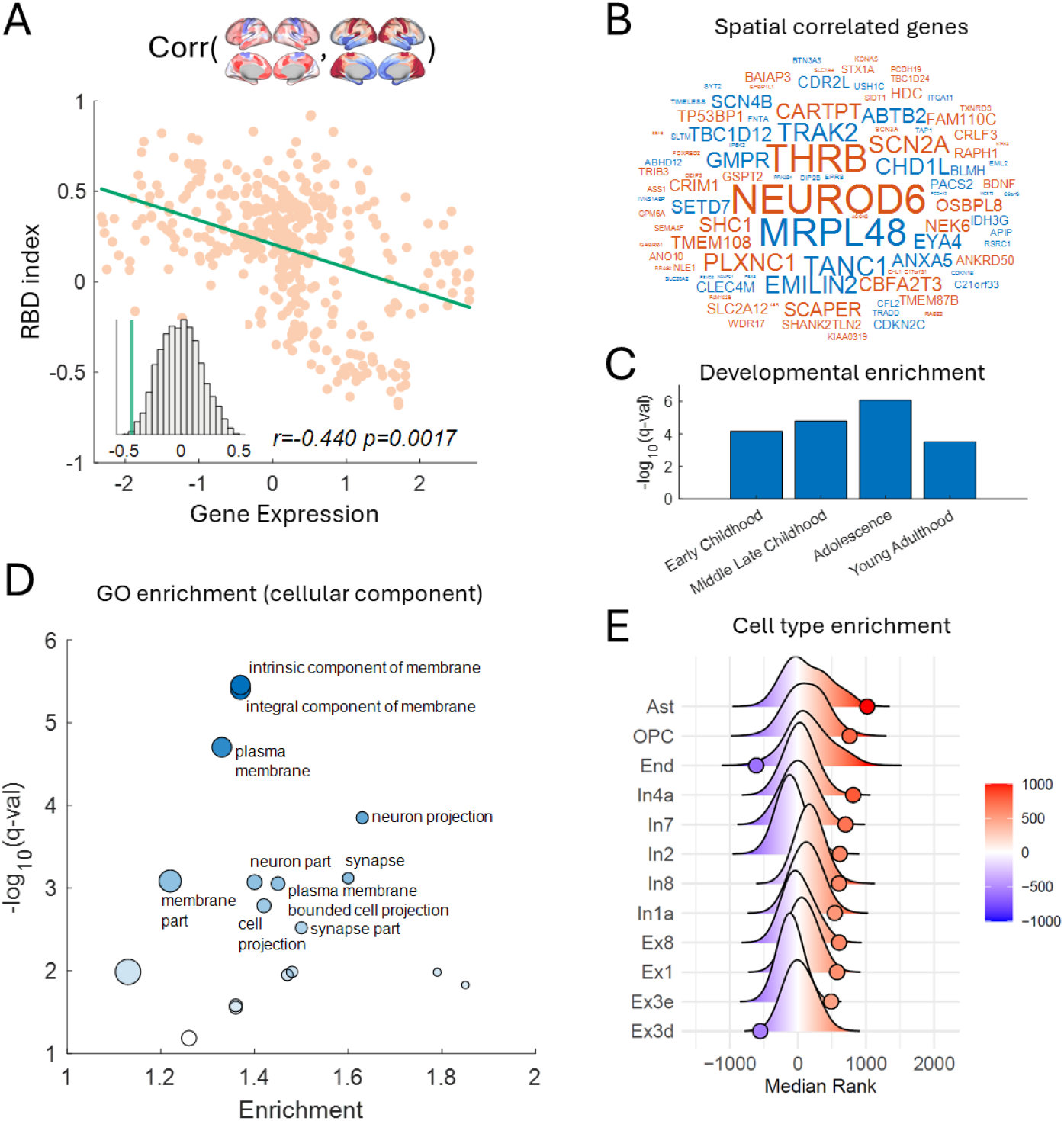
Associations between the regional FC development pattern of P2 and gene expression. (A) Alignment of P2 with gene expression PC1. (B) The top 100 genes with the highest absolute correlations with P2 are displayed; warm and cool colors indicate positive and negative correlations respectively, and font size is proportional to the magnitude of the absolute correlation. (C) Neurodevelopmental gene enrichment analysis revealed enrichment for the highly correlated neurodevelopmental genes expressed in the cortex during developmental stages ranging from early childhood to young adulthood. The y-axis represents the statistical significance of the enrichment (FDR-corrected *q*-value). (D) Gene Ontology (GO) enrichment analysis for cellular components identified significant enrichment in terms such as “membrane,” “neuron projection,” and “synapse.” Each dot represents an enriched GO term, with size proportional to the number of associated genes and transparency reflecting the significance of enrichment (FDR-corrected *q*-value, y-axis). (E) Adult cell-type-specific enrichment analysis demonstrated that the correlated genes were enriched for markers of various cortical cell types in adults, including astrocytes, oligodendrocyte precursor cells (OPCs), and multiple inhibitory and excitatory cortical neurons.

We further performed Gene Ontology (GO) enrichment analysis on the set of correlated genes using the online tool of GOrilla [36], which revealed that the significantly correlated genes were primarily enriched in GO terms associated with “membrane”, “neuron projection”, and “synapse” (*q*<0.05, FDR corrected, Fig. 4D). Given that regional differences in cortical gene expression may be influenced by differences in cellular composition of the cortex, we conducted a cell-type-specific enrichment analysis [37] to examine the enrichment of correlated genes in adult cortical cell types. This analysis revealed that the correlated genes were enriched across multiple adult cell types, including astrocytes, oligodendrocyte precursor cells, inhibitory and excitatory neurons (Fig. 4E).

### The FC development patterns are generalizable to the HCP-Development cohort

To assess the robustness of our findings, we investigated regional FC development using data from the HCP-D cohort (*n*=456; ages 5-22) and evaluated whether the identified FC development patterns could stratify the HCP-D individuals into distinct subgroups. Five-fold cross-validation was utilized to predict brain age for all cortical regions and individuals in the HCP-D cohort, as in the previous analyses of the PNC cohort. The spatial variation of FC development across cortical regions exhibited a pattern consistent with that observed in the PNC cohort (Spin-test, *r*=0.467, *p*<0.0001), with regions showing higher predictive accuracy primarily located in the somatomotor network, VAN, and the anterior part of the DMN (Fig. 5A and 5B). Next, HCP-D individuals were stratified into three subgroups based on the FC development patterns identified in the PNC cohort. Specifically, each individual’s RBD index map was compared with the PNC cohort’s centroids of RBD index maps (P1, P2, and P3), and the individual was assigned to the closet matching subgroup (Fig. 5C). We then analyzed cognitive performance of individuals across subgroups in three domains: fluid cognition, crystallized cognition, and total cognition. Individuals in subgroup P2 demonstrated the highest cognitive performance, with significant differences in fluid cognition and total cognition compared to subgroup P1 (Wilcoxon rank sum test, *p*<0.05, Fig. 5D). As in the PNC cohort, no significant differences were observed between subgroups P1 and P3. These findings demonstrate that the three FC development patterns identified in the PNC cohort are robust and generalizable to the HCP-D cohort. Moreover, the results reinforce that the spatial heterogeneity in FC development across individuals is linked to neurocognitive performance.

**Fig 5.**
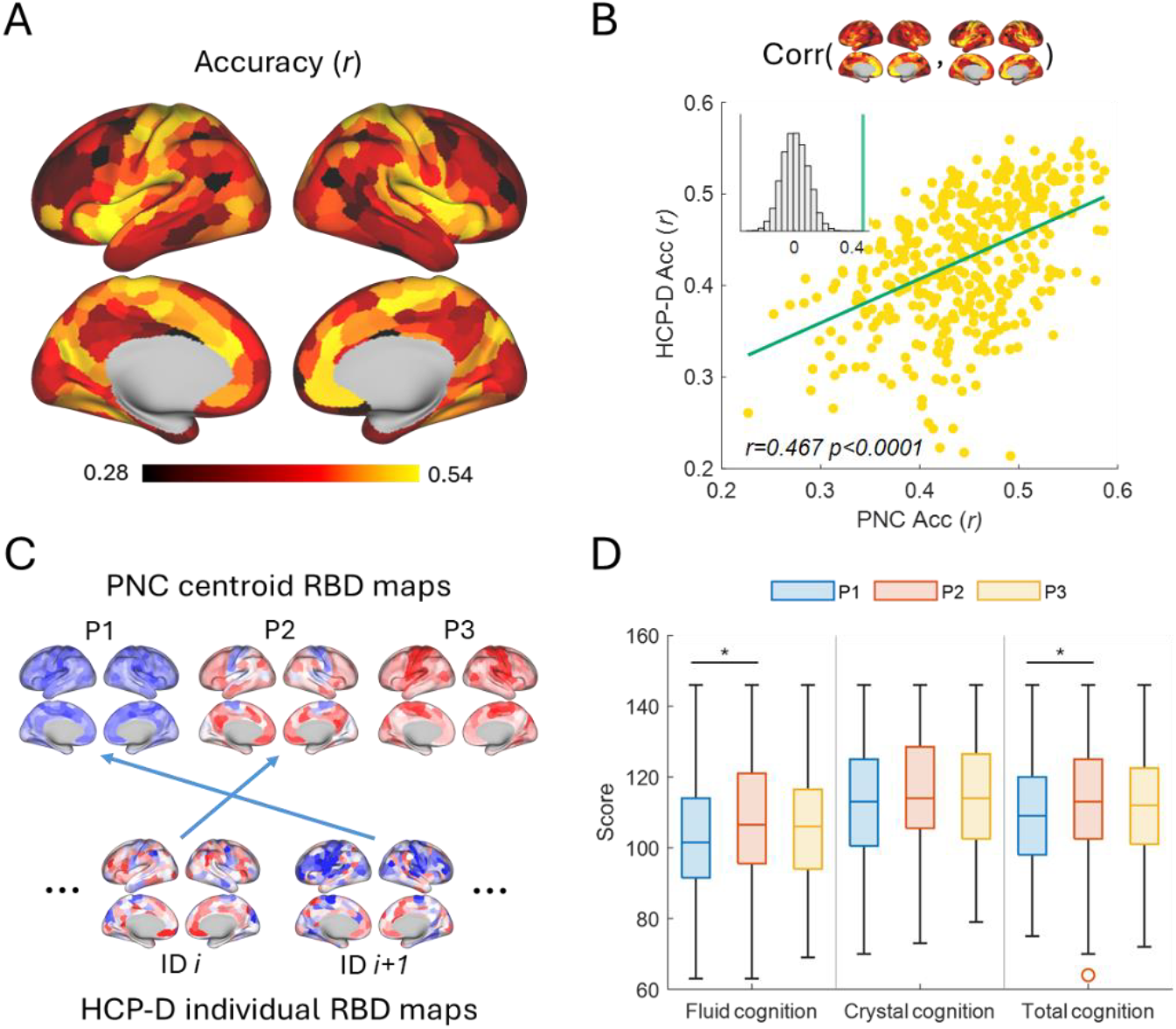
Replication analysis on the HCP-Development cohort. (A) Regional capacity for age prediction, measured by accuracy (*r*). (B) Spatial alignment of the PNC accuracy (*r*) with that of HCP-D, revealed consistent pattens of regional FC development in both cohorts (*p*<0.0001, Spin-test). (C) Assignment of HCP-D individuals to the FC development subtypes defined in the PNC cohort. For each HCP-D individual, the distances between its RBD index map and the centroid RBD index maps of the PNC-defined subtypes were calculated, and the individual was assigned to the subtype with the smallest distance. For example, individuals *i* and *i+1* were assigned to subtypes P2 and P1, respectively. (D) Statistical comparisons of cognition scores across three subgroups in HCP-D, highlighting subtype-related differences in cognitive profiles.

## Discussion

In this study, we investigated the spatial heterogeneity of brain FC development at the individual level by leveraging machine learning-based brain age prediction and fMRI data of large cohorts of youths. Using a region-specific brain age prediction approach, we estimated an RBD index for each brain region and individual, uncovering significant variability in FC development across cortical regions, which reflects the spatial heterogeneity of brain FC development at the individual level. By clustering these individualized RBD index maps, we identified three distinct development patterns and stratified individuals into corresponding subgroups. Notably, individuals with a FC development pattern positively correlated with the hierarchical cortical organization exhibited superior cognitive performance, a relationship that could not be captured using global BAG measures. These findings underscore the utility of RBD indices in providing fine-grained insights into the complex relationships between brain development and behavior phenotypes, offering a more nuanced understanding than global measures can achieve.

Previous research has investigated brain development in children and adolescents using imaging-based brain age estimation. Multimodal brain development indices have revealed that elevated brain age is associated with significantly lower gray matter volume, higher white matter volume in the prefrontal cortex, and enhanced FC in networks such as the DMN [9]. Additionally, brain age estimates have been linked to various cognitive functions [5] and psychiatric disorders, including schizophrenia [9], autism [11], and attention-deficit/hyperactivity disorder [13]. However, most of these studies focus on brain development from a global perspective by estimating BAG, a single univariate index, for each individual. This approach overlooks the spatial heterogeneity of brain development, which reflects the possibility that brain regions develop at varying rates within the same individual. While cluster-wise brain development indices derived from structural imaging data are promising to capture brain changes that are not detectable by BAG [14], the region-wise heterogeneity of FC development within individuals remains largely unexplored. This study systematically characterizes the individualized RBD index of FC in children and adolescents, providing new insights into the spatial heterogeneity of FC development. Our findings reveal that this heterogeneity reflects differences in developmental pacing across brain regions and is significantly linked to individual differences in cognitive performance.

We observed spatial heterogeneity in FC development at both the population and individual levels. At the population level, different brain regions exhibited varying capacities for age prediction, with regions within the somatomotor network and VAN showing higher prediction accuracy. At the individual level, brain regions displayed heterogenous advanced or delayed brain development across cortical regions, independent of an individual’s chronological age. The RBD indices facilitated a fine-grained, individualized characterization of the FC development, enabling the stratification of individuals into subgroups based on distinct FC development patterns. While it might seem intuitive that individuals with advanced FC development would outperform those with overall delayed development in cognitive tasks, our findings highlight the critical role of specific brain regions in this relationship: individuals with advanced development in the DMN and VAN regions demonstrated superior cognitive performance, whereas those with advanced development in the somatomotor regions did not, despite showing greater overall advancement in FC development. These results underscore the importance of accounting for spatial heterogeneity in brain age-related studies, providing compelling evidence that region-specific patterns of development significantly influence cognitive outcomes.

The subtypes of FC development identified by the global BAG exhibited distinct cognitive profiles compared to those derived based on the RBD index maps. Notably, its subgroup with the best cognitive profile corresponded to the subgroup in-between the delayed and advanced developmental groups (Fig. S1A). Similarly, a recent study found that the global BAG had limited utility in capturing fluid cognition in older individuals [18]. Our findings further reveal that the distributions of global BAG measures were largely overlapping across subgroups identified based on the RBD indices (Fig. S1B), indicating that the RBD indices provides additional insights into brain development beyond what is captured by global brain development index alone. These results highlight the potential of incorporating the spatial heterogeneity in brain age to better capture the associations between brain age and neurocognition.

The regional brain development pattern of the subgroup with superior cognitive performance aligns significantly with the brain’s hierarchical organization. Advanced development predominantly occurs in regions with high hierarchy ranks, such as DMN and VAN regions, and delayed development occurs in regions with low hierarchy ranks, such as sensorimotor regions. This finding is consistent with previous research showing that the development of hierarchical functional system is supportive to neurocognitive development [1, 2, 38]. It also corroborates findings that individual FC patterns in the DMN and VAN are linked to cognitive functions in youth [20-22]. Furthermore, the regional brain development pattern shows a significant correlation with cortical myelination, suggesting that changes in cortical microstructure may influence FC development by enhancing neural signal transmission and regulating plasticity [39]. These associations converge to support the idea that advanced FC development parallel to the hierarchical axis of cortical organization underpins better cognition in youth.

We further investigated the biological basis of the advanced FC development pattern through gene expression data analysis, identifying a set of co-located genes, including NEUROD6 and THRB, known to play critical roles in brain development and neurocognition. These genes regulate and influence key processes such as neural cell migration and differentiation, synaptogenesis, and myelination [40-44]. Additionally, these co-located genes were enriched for those expressed in cortical regions during mid-to-late childhood and adolescence, consistent with known developmental trajectories [35]. Gene ontology enrichment analysis revealed that these genes were associated with multiple cellular components, including membrane, neuron projection, and synapse. Cell-type-specific enrichment analysis further demonstrated correlations between the advanced FC development pattern and genes related to astrocytes as well as multiple excitatory and inhibitory neuron types. These findings suggest that the co-located genes play essential roles in supporting the inter-neuron connectivity and brain circuit development [45]. It is speculated that the identified FC development pattern reflects underlying changes in gene transcription and its influence on brain cytoarchitecture.

We also observed that the spatial heterogeneity of FC development was both consistent and replicable across different datasets. Predictive analysis on the HCP-D cohort demonstrated that the region-wise predictive capacity closely aligned with that observed in the PNC cohort. Additionally, the FC development patterns identified in the PNC cohort were generalizable to the HCP-D cohort. Stratification results from the HCP-D cohort demonstrated that individuals in the subgroup P2 exhibited superior cognitive performance. Notably, this was observed despite differences in the cognition domains assessed in the PNC and HCP-D cohorts, indicating that the identified FC development patterns were robust and reliably captured underlying brain development processes.

Although the RBD index has demonstrated potential for capturing fine-grained and reproducible patterns of brain development, several limitations should be acknowledged. First, though brain age has been validated to be sensitive to clinical outcomes and other biomarkers [46-48], concerns have been raised regarding the inflated prediction accuracy due to statistical bias-correction and its interpretation [49]. While brain age prediction accuracy is not the focus of this study, alternative measures merit further investigation. Second, the use of a linear modeling approach for region-wise brain age prediction may constrain its ability to characterize non-linear developmental effects in certain brain regions. Developing a principled framework to select the optimal predictive model (linear or non-linear) for each region could better characterize the complex developmental changes in FC. Third, the associations between FC development patterns and gene expression data rely on the Allen Human Brain Atlas, which represents transcriptomic data from adult brains. Further replication studies are needed when transcriptome data from the youth population becomes available, to confirm and extend these findings. Fourth, while sex was treated as a covariate and controlled in this study, investigating sex differences in the spatial heterogeneity of brain FC development offers a compelling avenue for future research. Lastly, this study focuses on FC development in youth, but integrating multi-modal imaging data and investigating the heterogeneity of brain development from a multi-dimensional perspective presents a promising direction for future investigations. Such approaches could provide a more comprehensive understanding of the complex mechanisms underlying brain development.

In summary, we investigated the spatially fine-grained development of FC in youth using the RBD index, derived by region-specific brain age predictive modeling. Our findings reveal replicable spatial heterogeneity in FC development at the individual level and establish links between the spatial patterns of region-wise brain development and neurocognition. Additionally, these results suggest region-wise brain development patterns align with the brain’s hierarchical organization and are potentially associated with gene expression. These findings underscore the critical role of spatial heterogeneity in FC development for supporting neurocognition development in youth.

## Methods

### Participants and data processing

Data were drawn from two datasets: the Philadelphia Neurodevelopmental Cohort (PNC; *n*=693) and the Human Connectome Project Development (HCP-D; *n*=456).

The PNC [27] is a community sample of children and adolescents from the greater Philadelphia area, collected for studying brain development, including 1601 participants with neuroimaging data. For this study, demographic, cognitive assessment, and neuroimaging data from 693 participants (301 males and 392 females) aged 8–23 years were included after applying the inclusion criteria as previously described [8]. Informed consent was obtained from all participants or their parents/guardians, and minors provided assent. All study procedures were approved by the institutional review boards of both the University of Pennsylvania and the Children’s Hospital of Philadelphia. One resting-state and two task-based (n-back and emotion identification) fMRI scans for each participant were used in this study. The neuroimaging preprocessing followed previously described procedures and methods [8]. Cognitive assessment was conducted using the Penn computerized neurocognitive battery (Penn CNB), and three factor-analysis based composite scores [50], including executive and complex cognition, episodic memory, and social cognition, were included for analysis in this study.

The HCP-D [28] includes 652 participants recruited across four sites in the U.S. to study healthy brain development in children and adolescents. A sample of 456 participants (214 males and 242 females) aged 5–22 years with complete demographic, cognitive assessment, and neuroimaging data were included in this study after applying the inclusion criteria as previously described [3]. All study procedures were approved by a central Institutional Review Board at Washington University in St. Louis. Resting-state and task-based (carit, emotion, and guessing tasks) fMRI scans were used for all participants. The neuroimaging acquisition and preprocessing procedures have been previously described [3]. Participants completed a battery of cognitive assessments, including segments of the NIH toolbox and others [28]. Composite scores including fluid cognition, crystalized cognition and total cognition from the HCP-D data release [28] were included in this study.

For both samples, preprocessed fMRI time series were parcellated into 400 cortical regions using the Schaefer-400 atlas [51], and time series from resting-state and task fMRI scans were concatenated to compute FC measures for the analysis.

### Brain development index

For region-specific brain age prediction, a dedicated linear ridge regression model to predict chronological age was built for each cortical region on its FC profile that was defined as its FC values to all the other cortical regions, with each value measured as a Pearson’s correlation coefficient between the two regions. Five-fold cross-validation was utilized to obtain predicted brain age for all individuals and cortical regions.

For global brain age prediction, the whole-brain functional connectome (upper triangle entries of the FC matrix, containing FC values between all pairs of cortical regions) was used as features, and a linear ridge regression model was trained for the prediction of chronological age.

Both the region-specific and global prediction models were built and evaluated with the same five-fold cross-validation, with identical training and evaluation data splits. For each prediction model, the regularization parameter in ridge regression was optimized in the range [2^™10^, 2^™9^, …, 2^4^, 2^5^] using nested cross-validation within the training sample.

Given the predicted brain age for each cortical region, the difference between this measure and chronological age was referred as the regional brain development (RBD) index, reflecting region-specific FC development. To alleviate the known systematic bias in the brain age prediction due to the “regression to the mean” effect, the RBD index was adjusted by a linear bias correction procedure [30]. In addition, covariates including sex, head motion, and site information (for the HCP-D sample) were regressed out from the RBD index for all subsequent analysis. The same procedure was applied to obtain global brain age gap (BAG) measures based on the predicted global brain age.

### Subtyping FC development

Given the RBD index maps for all participants of the PNC cohort, the K-means clustering algorithm was utilized to identify subtypes of FC development and stratify the participants into different subgroups. Specifically, the clustering was performed with whole-brain RBD indices as features and cosine distance as the distance metric, as implemented in MATLAB (R2021a).

The number of clusters was set to 3, determined based on both reproducibility and within-cluster point-to-centroid distances of the clustering results. Specifically, all participants were randomly split into two halves, and clustering was performed on each half separately with the number of clusters *k* set from 2 to 7. This procedure was repeated for 100 times. The clustering results for each half was denoted as *C*_*k*,1_ and *C*_*k*,2_. For each *k*, the clustering centroids of the first half were then applied to samples in the second half to obtain the clustering results *C*_*k*,1→2_, and vice versa to obtain *C*_*k*,2→1_. The average of *RandIndex*(*C*_*k*,1_, *C*_*k*,2→1_) and *RandIndex*(*C*_*k*,2_, *C*_*k*,1→2_) was adopted as the reproducibility measure for clustering results with *k* clusters, where *RandIndex*(*C,C*^′^) refers to the Rand index for two sets of clustering results *C* and *C*^′^. Additionally, to evaluate the data-fitting performance of the clustering results, the distance of each data point to its corresponding cluster centroid was calculated and averaged across all data points. The optimal number of clusters was determined by reaching a balance between reproducibility and data-fitting performance, where the two measures interacted as shown in Fig. S4.

The HCP-D individuals were stratified into distinctive subgroups based on the FC development patterns identified in the PNC cohort. Specifically, each individual’s RBD index map was compared to the PNC cohort’s centroid maps, and the individual was assigned to the subgroup with the closet match. For instance, individuals *i* and *i+1* were assigned to subgroups P2 and P1, respectively, as their RBD index maps showed the highest similarity to the corresponding centroids (Fig. 5C).

### Meta-analytic functional decoding of FC development pattern

The meta-analytic decoding tool [31] was used to link the FC development pattern to functional/behavioral profiles. Specifically, the functional terms and term-based meta-analytic maps from the NeuroQuery database [32] was utilized with the correlation decoder [31] for the functional decoding. The functional decoding results was illustrated by a word cloud, with larger font size indicating higher functional relevance.

### Gene enrichment analysis

Gene expression data was obtained from the Allen Human Brain Atlas (AHBA) [33] and processed with the abagen toolbox [52] and the Schaefer-400 atlas in the MNI space to obtain regional gene expression. Since not all genes in the AHBA dataset are assumed to be expressed in the brain, only brain-expressed genes were retained for analysis, following an established procedure [53]. Additionally, the analysis focused on the left hemisphere with 200 brain regions to minimize sample variability across brain regions, as the gene expression data were sampled from both hemispheres of only two of the six donors [54]. A total of 4112 genes were included in our analysis.

The spatial alignment of regional FC development patterns and the expression patterns of each gene was quantified using Peason’s correlation coefficient, and its significance was tested using the spin test with 10000 permutations [55, 56]. The set of significantly correlated genes was included in the subsequent enrichment analysis. A Specific Expression Analysis (SEA) tool [35] was first used to examine the enrichment for cortical-expressed genes at different developmental stages through childhood to young adulthood. A rank-based Gene Ontology (GO) enrichment analysis using Gorilla [36] was then conducted to test the enrichment for the ontology of cellular component. As the regional differences in gene expression may reflect underlying differences in cellular composition across cortical regions, a median rank-based approach [53] was adopted to assess the enrichment for the adult cell types [37].

### Cortical organization maps

The sensorimotor-association axis [1] and the first principal component of gene expression [34] used in this study are available at https://github.com/PennLINC/S-A_ArchetypalAxis. The principal functional gradient map [57] and the myelin content map [58] are available at https://github.com/PennLINC/Brain_Organization.

## Data and code availability

The data used in this study are publicly available. The Philadelphia Neurodevelopmental Cohort is accessible from the Database of Genotypes and Phenotypes (phs000607.v3.p2) at https://www.ncbi.nlm.nih.gov/projects/gap/cgi-bin/study.cgi?study_id=phs000607.v3.p2. Data from the Human Connectome Project-Development are available for download through the NIMH Data Archive (https://nda.nih.gov/). Codes used for data analysis will be available at: https://github.com/MLDataAnalytics.

## Acknowledgements

This study was supported by grants from the National Institute of Health: R01EB022573, R01AG066650, U24NS130411.

## Competing interests

All authors declare no competing interests.

## Notes

### Competing Interest Statement

The authors have declared no competing interest.

