## Supplemental materials for "Spatial Heterogeneity and Subtypes of Functional Connectivity Development in Youth"

[1]Center for Biomedical Image Computing and Analytics, University of Pennsylvania, Philadelphia, PA 19104, USA
[2]Department of Radiology, University of Pennsylvania, Philadelphia, PA 19104, USA
[3]Chinese Institute for Brain Research, Beijing, 102206, China
[4]Department of Psychiatry, University of Pennsylvania, Philadelphia, PA 19104, USA
[5]Penn/CHOP Lifespan Brain Institute, University of Pennsylvania, Philadelphia, PA 19104, USA
[6]Department of Neurology, University of Pennsylvania, Philadelphia, PA 19104, USA
[7]Penn Statistics in Imaging and Visualization Center, University of Pennsylvania, Philadelphia, PA 19104, USA
[8]Department of Biostatistics, Epidemiology and Informatics, University of Pennsylvania, Philadelphia, PA 19104, USA
[9]Center for Brain Imaging and Stimulation, University of Pennsylvania, Philadelphia, PA 19104, USA
[10]Penn Brain Science, Translation, Innovation, and Modulation Center, University of Pennsylvania, Philadelphia, PA 19104, USA
[11]Center for Neuromodulation in Depression and Stress, University of Pennsylvania, Philadelphia, PA 19104, USA
[12]Departments of Neuroscience, Neurosurgery and Bioengineering, University of Pennsylvania, Philadelphia, PA 19104, USA
[13]Penn Lifespan Informatics and Neuroimaging Center, University of Pennsylvania, Philadelphia, PA 19104, USA

### Supplemental materials

#### Global brain age gap of distinct FC development subtypes

The global brain age gap (BAG) measure for each individual was calculated using a brain age prediction model trained on the whole-brain FC measures as features, employing the same 5-fold cross-validation approach used in the region-wise brain age predictive modeling.

Based on the global BAG measures that were corrected for bias and covariates with the same method used for the RBD indices, individuals were also stratified into three subgroups regarding global FC development: P1-G, P2-G, and P3-G, representing delayed, normative, and advanced development respectively. These subtypes exhibited different cognitive profiles compared with subtypes derived from RBD index maps. Specifically, individuals in P2-G exhibited significantly higher executive function scores than those in P1-G, while both P2-G and P3-G showed significantly higher social cognition scores than P1-G. However, no significant differences in memory scores were observed among the three subgroups (Fig. S1A).

The distributions of global BAG measures among the subgroups identified based on the RBD indices (P1, P2, and P3) showed considerable overlap (Fig. S1B), especially between subtypes P2 and P3. These results indicate that the patterns derived from the RBD indices are different from those based on the global BAG measures.

Together, these findings suggest that the RBD index maps, with its spatial heterogeneity, would capture fine-grained characteristics about the brain FC development, as different brain regions contribute to distinct functional roles. In contrast, the global BAG measures may obscure the heterogeneity of regional development by aggregating it into a single measure.

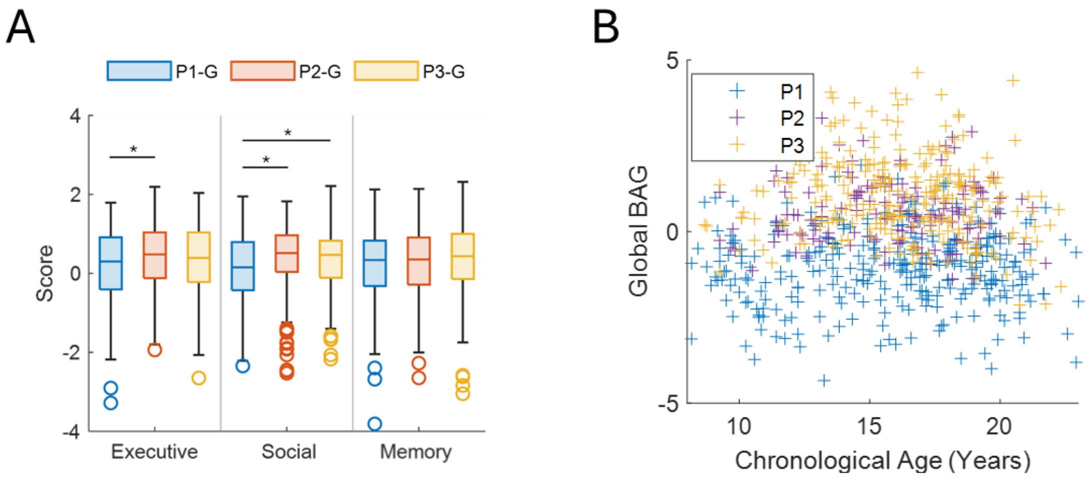


**Fig. S1**. Global brain age gap (BAG) of distinct FC development subtypes. (A) Cognitive scores of individuals across three subtypes defined by the global BAG measures (P1-G, P2-G, and P3-G). (B) Scatter plot of global BAG measures, color-coded by the subtypes derived from the RBD indices (P1, P2, and P3). Each '+' marker represents an individual.

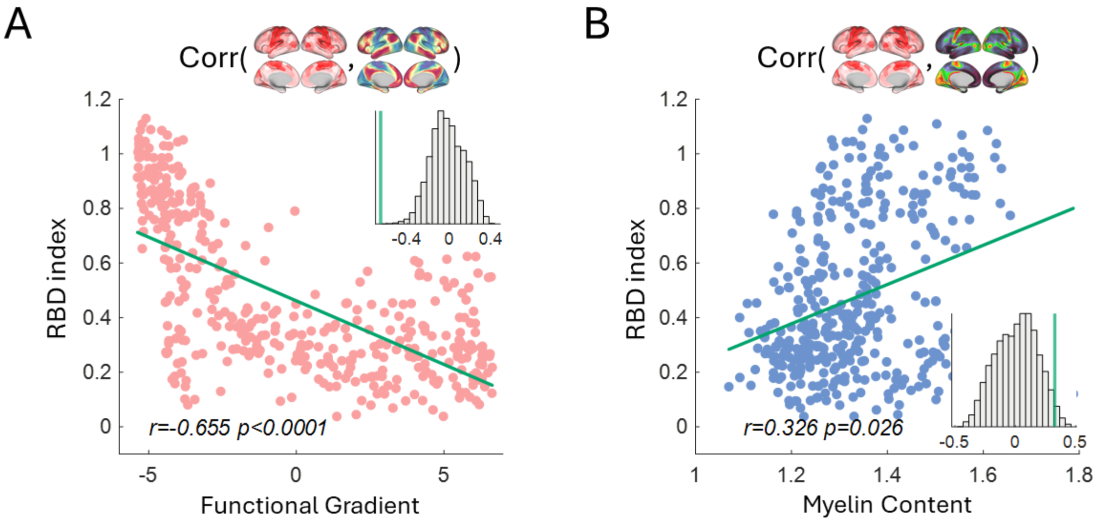


**Fig. S2**. (A) Significant negative correlation between the development pattern P3 and the principal functional gradient ($p$<0.0001, Spin-test). (B) Significant correlation between the development pattern P3 and the cortical myelin content ($p$=0.026, Spin-test).

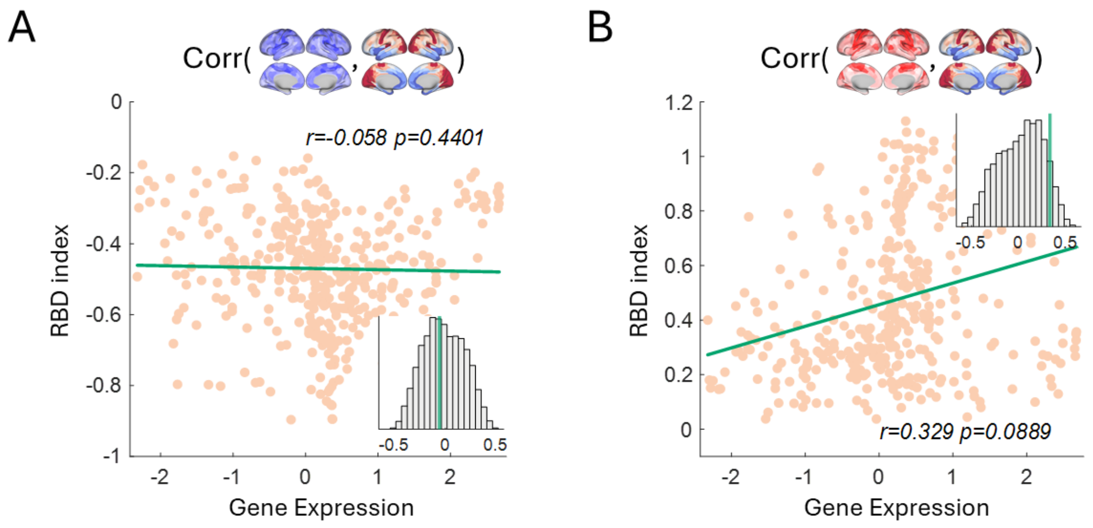


**Fig. S3**. No significant associations of the development patterns P1 and P3 with the first principal component of gene expression ($p$>0.05, Spin-test).

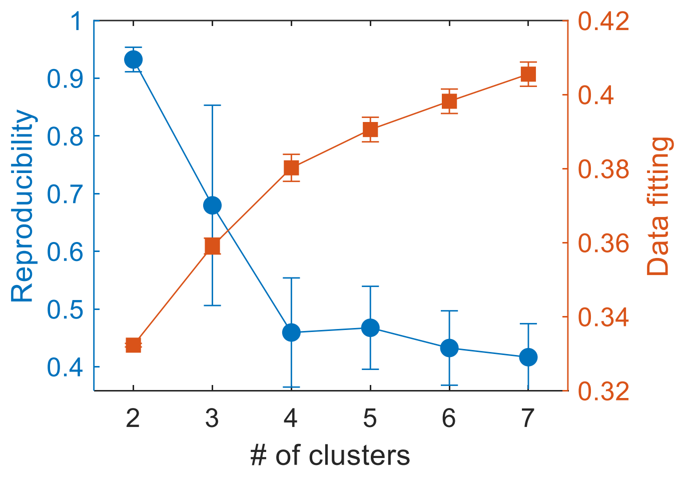


**Fig. S4**. Reproducibility and data-fitting measures for FC development subtyping across different numbers of subtypes (2 to 7) over 100 split-half replications. The Rand index was used as the reproducibility measure, and the data-fitting measure was computed as $1 - d_{avg}$, where $d_{avg}$ was calculated as the average of all data points' distances to their corresponding cluster centroids.
